# A single low-dimensional neural component of motor unit activity explains force generation across repetitive isometric tasks

**DOI:** 10.1101/2025.02.04.635938

**Authors:** Hélio V. Cabral, J. Greig Inglis, Elmira Pourreza, Milena A. Dos Santos, Caterina Cosentino, David O’Reilly, Ioannis Delis, Francesco Negro

**Author notes:** <u>Corresponding author</u>: Prof. Francesco Negro, Department of Clinical and Experimental Sciences, Università degli Studi di Brescia, Viale Europa 11, Brescia, 25121, Italy.

## Abstract

Previous studies suggest that low-dimensional control underlies motor unit activity, with low-frequency oscillations in common synaptic inputs serving as the primary determinant of muscle force production. In this study, we used principal component analysis (PCA) and factor analysis (FA) to investigate the relationship between low-dimensional motor unit components and force oscillations during repetitive isometric tasks with similar force profiles. We assessed the consistency of these components across trials in both individual (tibialis anterior; first dorsal interosseous) and synergistic muscles (vastus medialis, VM; vastus lateralis, VL). Participants performed 15 trials of a force-matching learning task. Three post-skill acquisition trials were selected for analysis to ensure high similarity in force profiles. Motor units were decomposed from high-density surface electromyograms, tracked across trials, and their smoothed discharge rates were decomposed into low-dimensional components using PCA and FA. Parallel analysis indicated that a single component could explain the smoothed discharge rates for the individual muscles and two components for VM-VL. Importantly, the first component explained most of the variance (∼70%) in smoothed discharge rates across all muscles. The first motor unit component also showed significantly higher correlations with force oscillations than the second component and remained highly consistent across trials. These findings were further supported by a non-linear framework combining network- and information-theoretic tools, which revealed high motor unit network density in the first component of all muscles. Collectively, these results suggest that, during isometric contractions, motor unit activity is primarily controlled by a single dominant shared synaptic input that closely mirrors force oscillations.

## Introduction

The central nervous system (CNS) generates and controls movement by transmitting and modulating neural commands to muscles, ensuring precise coordination and force production (Liddell and Sherrington, 1925; Sherrington, 1925). Although movement execution often appears effortless, even simple and repetitive tasks, such as reaching or walking, involve remarkable complexity. These actions require the simultaneous activation of multiple muscles, each comprising thousands of motor units, to generate torque across multiple joints and produce purposeful motion (Bernshteĭn, 1967; Lacquaniti et al., 2012; D’avella and Lacquaniti, 2013). At every level of the motor control hierarchy, the CNS must coordinate a vast number of interacting elements while managing the hierarchical mismatch between their abundance (e.g., much greater number of alpha motor neurons than the number of muscles). While this redundancy in the neuromuscular system offers flexibility and adaptability, it also presents a complex control problem, as the CNS must navigate infinite potential solutions to achieve a given motor goal (commonly referred to as the "degrees of freedom problem"; Bernshteĭn (1967)).

Several theoretical frameworks have addressed the degrees of freedom problem (Latash et al., 2007; Bruton and O’Dwyer, 2018). Among them, a prominent hypothesis is that the CNS employs a modular control strategy rather than individually controlling each element, reducing the complexity and dimensionality of control (Tresch et al., 2002; Gentner and Classen, 2006; Giszter et al., 2007; Bizzi et al., 2008; Tresch and Jarc, 2009; Alessandro et al., 2013; Ting et al., 2015). One key formulation of this modular control is the muscle synergy hypothesis, which proposes that a repertoire of movements is generated by the coordinated activity of muscles, referred to as synergies, in weighted combinations (Tresch et al., 1999; d’Avella et al., 2003; Ting and Macpherson, 2005). Empirical evidence supporting the existence of muscle synergies has been gathered from both animal (Tresch et al., 1999; Saltiel et al., 2001; d’Avella et al., 2003; Overduin et al., 2008) and human (Soechting and Lacquaniti, 1989; Ivanenko et al., 2005; d’Avella et al., 2011) studies, where the observed muscle activity or kinematic patterns were modeled as linear combinations of a small set of components (Tresch et al., 2006). Despite its valuable contributions to understanding neuromuscular control, this body of research has primarily analyzed the muscles rather than the individual motor units activating them.

Recent advances in technology and analytical methods have shifted the scale of analysis from muscles to individual neurons. These developments have enabled the recording and analysis of the activity of several motor neurons at both cortical (Kipke et al., 2008; Cunningham and Yu, 2014; Gao and Ganguli, 2015; Steinmetz et al., 2021) and spinal (Holobar and Zazula, 2007; Muceli et al., 2015; Chen and Zhou, 2016; Negro et al., 2016b; Muceli et al., 2022) levels, significantly advancing the understanding of the dimensionality of neural control. At the cortical level, compelling evidence has shown that movement planning and execution are governed by a small set of neural activity patterns spanning a low-dimensional space, often referred to as the “neural manifold” (Gallego et al., 2017). Corroborating earlier research that investigated the collective behavior of neurons (De N, 1938; Hebb, 1949), this evidence supports the idea that the population, rather than single neuron, is the fundamental unit underlying neural dynamics (Saxena and Cunningham, 2019; Ebitz and Hayden, 2021; Yuste et al., 2024). A similar framework has been proposed at the spinal level, where examining the pool of motor units offers deeper insights into muscle force control (Farina and Negro, 2015). Simulation and experimental studies have extensively demonstrated that the alpha motor neuron pool acts as a highly selective filter, linearly transmitting the common synaptic inputs while attenuating independent inputs (Negro et al., 2009; Negro and Farina, 2011; Farina et al., 2014; Negro et al., 2016a; Thompson et al., 2018). These findings suggest that common synaptic inputs are the primary determinant of generated muscle force during isometric contractions, supporting the hypothesis that low-dimensional neural control extends to motor unit activity (Negro et al., 2009; Laine et al., 2015; Madarshahian et al., 2021; Del Vecchio et al., 2023; Hug et al., 2023; Levine et al., 2023; Rossato et al., 2024).

A defining feature of low-dimensional neural control is the coupling among elements within the system, which results in a high degree of correlation among neural outputs (De Luca et al., 1982; Georgopoulos et al., 1982; Farmer et al., 1993; Hatsopoulos et al., 1998). Consequently, dimensionality reduction techniques, such as principal component analysis (PCA) and factor analysis (FA), have been widely applied to discharge rates of neuronal ensembles to identify dominant patterns of covariation, yielding a reduced set of explanatory components that capture key control features (Cunningham and Yu, 2014). For instance, Negro et al. (2009) demonstrated that the first principal component, derived using PCA from the smoothed discharge rates of active motor units, explains oscillations in isometric force more effectively than the discharge rates of individual motor units. In general, there exists a strong justification for using PCA to investigate patterns of neuronal correlation (Chapin and Nicolelis, 1999; Negro et al., 2009; Churchland et al., 2010; Levine et al., 2023) since PCA is mathematically linked to Hebbian theory (Oja, 1989, 1992; Olshausen, 1998), one of the most experimentally validated theories of synaptic plasticity (Hebb, 1949). More recent studies using alternative rotations of low-dimensional components or other dimensionality reduction techniques have suggested that multiple components may underlie motor neuron activity, particularly when synergistic muscles are involved (Del Vecchio et al., 2023; Nuccio et al., 2024; Rossato et al., 2024). If low-dimensional control is indeed an effective strategy for simplifying neural control via common projections to spinal motor neurons, the resulting components should closely mirror force oscillations (i.e., the final motor output) and exhibit consistency across trials with similar force outputs.

In this study, we investigated the relationship between low-dimensional components of motor unit activity and oscillations in muscle force during repetitive isometric tasks with highly similar force outputs. We further examined the consistency of these components across trials both for individual (tibialis anterior, TA; first dorsal interosseous, FDI) and synergistic muscles (vastus medialis, VM; vastus lateralis, VL). Participants performed fifteen trials of a force-matching learning task, and, following the learning phase, three consecutive trials were analyzed to ensure high similarity in force oscillations. Motor units were decomposed from high-density surface electromyograms (HDsEMG), tracked across these trials, and their smoothed discharge rates were linearly decomposed into low-dimensional components using PCA and FA. To further characterize the coupling among motor unit activity across trials, we employed a recently developed non-linear framework combining information and network theoretic tools to decompose neural signals and characterize their network structure (O’Reilly and Delis, 2022, 2024). We hypothesized that a single low-dimensional component, highly resembling force oscillations, would show strong consistency across trials, regardless of the muscle analyzed or method employed.

## Methods

### Participants

Twenty-nine healthy participants performed a series of isometric force-matching tasks involving index finger abduction (FDI muscle), dorsiflexion (TA muscle), and knee extension (VM-VL muscles). Specifically, ten volunteers (4 females; age 31 ± 4 years; height 177 ± 8 cm; mass 72 ± 17 kg) participated in the FDI experiments, twelve volunteers (5 females; mean ± SD: age 31 ± 3 years; height 175 ± 9 cm; mass 69 ± 18 kg) in the TA experiment, and seven male volunteers (age 33 ± 10 years; height 184 ± 5 cm; mass 85 ± 19 kg) in the VM-VL experiment. Seven volunteers participated in both the TA and FDI experiments in separate sessions. All participants had no history of upper or lower limb injuries that could impact their ability to perform voluntary contractions. Before beginning the experiments, participants provided informed consent following an explanation of the experimental procedures. This study was approved by the local ethics committee of the University of Brescia (code NP5665) and conducted in accordance with the latest version of the Declaration of Helsinki.

### Experimental protocol

**Figure 1** illustrates the participants’ positioning for the index finger abduction, dorsiflexion, and knee extension tasks. For the index finger abduction task, participants were seated with their right wrist neutrally positioned on a custom-built device with their elbow flexed at 45° (0° being fully extended). The index finger was secured to an adjustable support attached to a load cell (SM-100 N, Interface, Arizona, USA) to record the isometric abduction force produced by the index finger (**Figure 1A**). To minimize the involvement of other muscles, the wrist was secured to the device with Velcro straps, and the other fingers (little, middle and ring) were strapped separately from the index finger. The thumb was fixed to an adjustable support, maintaining an approximate angle of 80° to the index finger (**Figure 1A**). For the dorsiflexion task, participants were seated with their right leg positioned on a custom-made ankle dynamometer. The knee was fully extended, the hip flexed at 70° (0° being fully extended), and the ankle joint at 10° of plantar flexion (0° being the foot perpendicular to the shank). The foot was secured with straps to a footplate connected to a load cell (SM-500 N, Interface, Arizona, USA) to measure the isometric dorsiflexion force (**Figure 1B**). To ensure that the force generated was exclusively due to the dorsiflexor muscles, additional straps were applied around the thigh and knee. For the knee extension task, participants were seated on an isokinetic dynamometer (Humac Norm Extremity System, CSMi Solutions, Massachusetts, USA) with their right knee flexed at 90° (0° being fully extended) and aligned as coaxially as possible with the dynamometer’s axis of rotation (**Figure 1C**). The hip was flexed at 90° (0° being fully extended), and the ankle joint was secured to the device approximately three centimeters above the malleolus, allowing for the measurement of isometric knee extension force. The trunk and waist were also secured to the device with Velcro straps.

**Figure 1:**
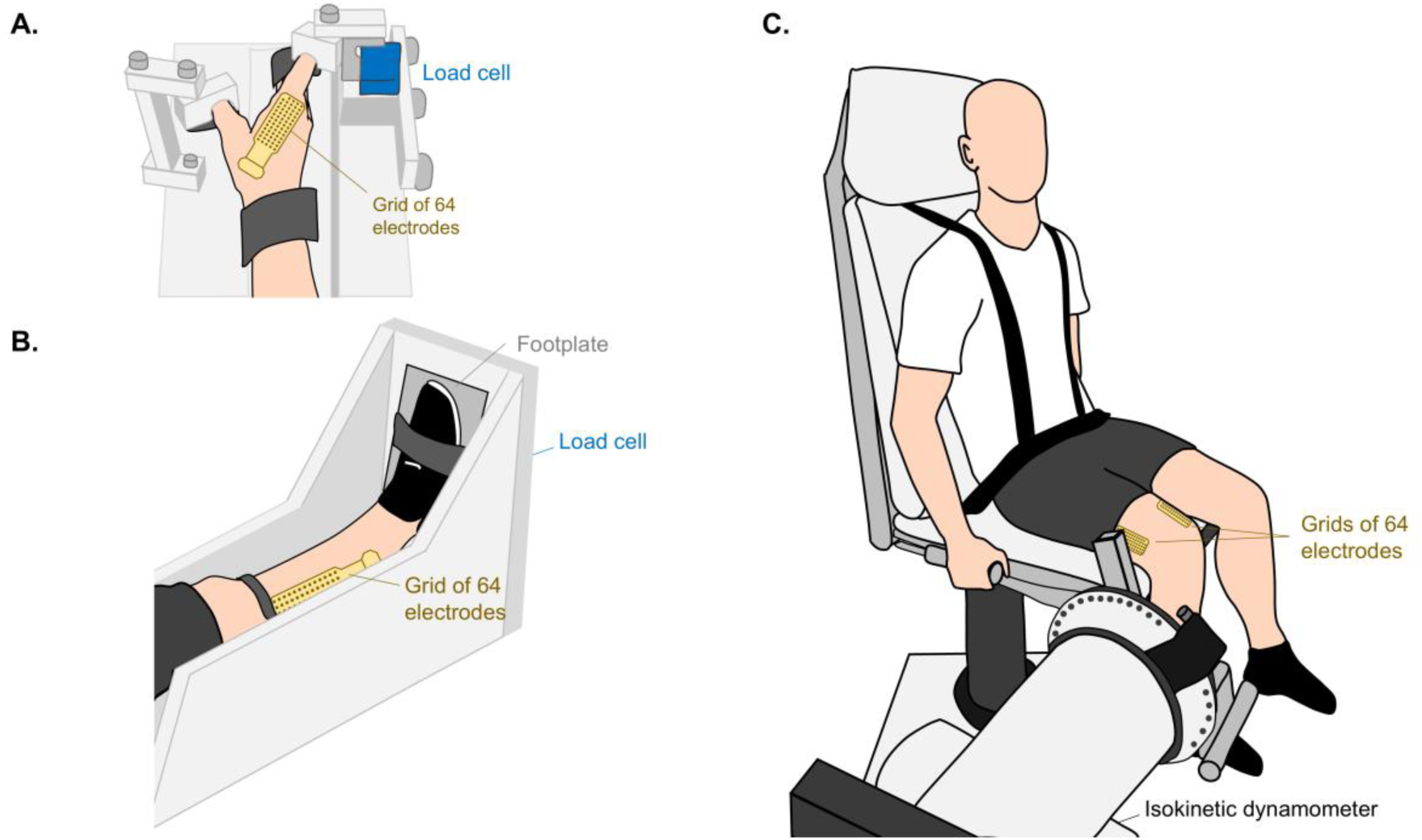
Experimental setup. Participants’ positioning for the index finger abduction (A), dorsiflexion (B), and knee extension (C) tasks. The isometric force produced by participants was measured using load cells (A, B) or an isokinetic dynamometer (C). High-density surface electromyography grids with 64 electrodes were used to record muscle activity from the first dorsal interosseous (A), tibialis anterior (B), and vastus medialis and vastus lateralis (C). For all tasks, participants performed an isometric force-matching task following a complex trajectory for 15 trials (learning task). The target force level was set at 5% of maximum voluntary contraction for the first dorsal interosseous and 10% MVC for the tibialis anterior and vastii muscles.

The experimental procedures were the same for the three muscle groups (FDI, TA, and VM-VL). Initially, participants performed three isometric maximal voluntary contractions (MVCs) for 3 s, with 2 min of rest between each contraction. The highest MVC among the three trials was used as a reference for the subsequent submaximal tasks. After a 5-min rest interval, participants were asked to perform an isometric force-matching task, following a complex trajectory for fifteen trials (learning task). This number of trials was chosen based on previous studies, which have demonstrated that fifteen trials of a similar task are sufficient for short-term learning (Knight and Kamen, 2004; Cabral et al., 2024b). The trajectory specifically involved a linear increase from 0% MVC to the target force level at a rate of 5% MVC/s, a stochastic force region at the target force level for 30 s, and a linear decrease from the target force level to 0% MVC at a rate of -5% MVC/s (Cabral et al., 2024b). The stochastic force region consisted of a randomly generated signal, low-pass filtered at 1.5 Hz, with oscillations around the target force level, set at 5% MVC for the FDI muscle and 10% MVC for the TA and VM-VL muscles. Three different trajectories were generated, one for each muscle group, but the same trajectories were used for all trials and participants. A rest period of at least 1 min was provided between trials. Throughout the task, participants were encouraged by the same researcher to match the trajectory as closely as possible. Visual feedback from the target and produced force was displayed on a computer monitor positioned approximately 60 cm in front of the participant.

### Data collection

Monopolar high-density surface electromyograms (HDsEMG) were collected from the TA, FDI, and VM-VL muscles during the submaximal isometric tasks using adhesive grids of 64 electrodes arranged into 13 rows by 5 columns, with a missing electrode in the upper left corner (OT Bioelettronica, Turin, Italy). For the TA and VM-VL muscles, grids with an 8 mm inter-electrode distance were used (GR08MM1305), while for the FDI muscle, grids with a 4 mm inter-electrode distance (GR04MM1305) were employed. The grids were positioned longitudinally along the muscle belly (**Figure 1**), which was identified through palpation by an experienced investigator. In the case of the VL muscles, two grids were used: one positioned distally and the other proximally. Before placing the electrodes, the skin was shaved and cleaned with abrasive paste (EVERI, Spes Medica, Genova, Italy) and water to enhance signal quality. Conductive paste (AC cream, Spes Medica, Genova, Italy) was applied in the foam cavities of the grid to ensure optimal electrode-skin contact. The reference electrode was positioned on the right ankle for TA and VM-VL muscles and on the right wrist for the FDI muscle. HDsEMG and force signals were sampled synchronously at 2048 Hz using a 16-bit A/D amplifier (10-500 Hz bandwidth; Quattrocento, OT Bioelettronica, Turin, Italy).

### Data analysis

All analyses were conducted offline using custom-written scripts in MATLAB (version 2022b; The MathWorks Inc., Natick, Massachusetts, USA).

#### Trial selection for analysis

Three consecutive trials were selected out of the fifteen performed, focusing on those after the learning of the force-matching skill (i.e., post-skill acquisition trials). Initially, the force signal was low pass filtered at 15 Hz using a third-order Butterworth filter. The root-mean-square error (RMSE) between the detrended force and target signals was then calculated considering the 30-s middle region of each trial. The three consecutive trials with the lowest RMSE between the force and target signals were selected for further analysis, as the force oscillations are expected to be most similar across these trials. For these selected trials, the coefficient of variation of force (i.e., standard deviation divided by mean) was calculated to quantify force steadiness.

#### HDsEMG decomposition and motor unit tracking

Due to the late recruitment of motor units, the first 5 seconds of data were discarded, and only the last 25 seconds of the middle region of the three selected trials were analyzed. Initially, monopolar HDsEMG signals were filtered between 20 and 500 Hz using a third-order bandpass Butterworth filter (**Figure 2A**). All signals were then visually inspected, and those of low quality (e.g., artifacts or skin-electrode problems during acquisition) were discarded from further analysis. The remaining HDsEMG signals were decomposed into motor unit discharge times using a convolutive blind-source separation algorithm (Negro et al., 2016b), which has been previously validated and widely applied to assess individual motor units in the muscles investigated in this study (Negro et al., 2016b; Martinez-Valdes et al., 2017; Cabral et al., 2024b). The identified motor units were visually inspected by an experienced operator, and missing or misidentified discharges (inter-spike intervals < 20 ms or > 250 ms; Negro et al. (2009)) were manually and iteratively edited (Martinez-Valdes et al., 2017; Hug et al., 2021). Motor units were then tracked across the three trials by reapplying the motor unit separation vectors, following procedures used in previous studies (Oliveira and Negro, 2021; Rossato et al., 2022; Cabral et al., 2024b). Briefly, the deconvolution of HDsEMG signals using independent component analysis involves the calculation of a separation matrix, whose columns are the separation vectors for each motor unit (Negro et al., 2016b; Dai and Hu, 2019). These separation vectors are unique for each motor unit and define the spatiotemporal filters that, when applied to the HDsEMG signals, yield the estimated motor unit discharge times. To maximize the number of tracked motor units, the estimated separation vectors from one trial were applied in the other two trials, considering all possible combinations. However, in a few cases (1 participant for TA, 3 for FDI and 2 for the VM), it was not possible to track more than one unit using this method. In such cases, motor units were tracked based on their action potential shapes (Martinez-Valdes et al., 2017). Specifically, the two-dimensional representation of the motor unit action potentials identified in one trial was estimated using the spike-triggered averaging technique and then cross-correlated with the spatial representation of the motor units identified in the other two trials. Motor units with high similarity between action potentials (cross-correlation > 0.8) were considered to belong to the same motor unit.

**Figure 2:**
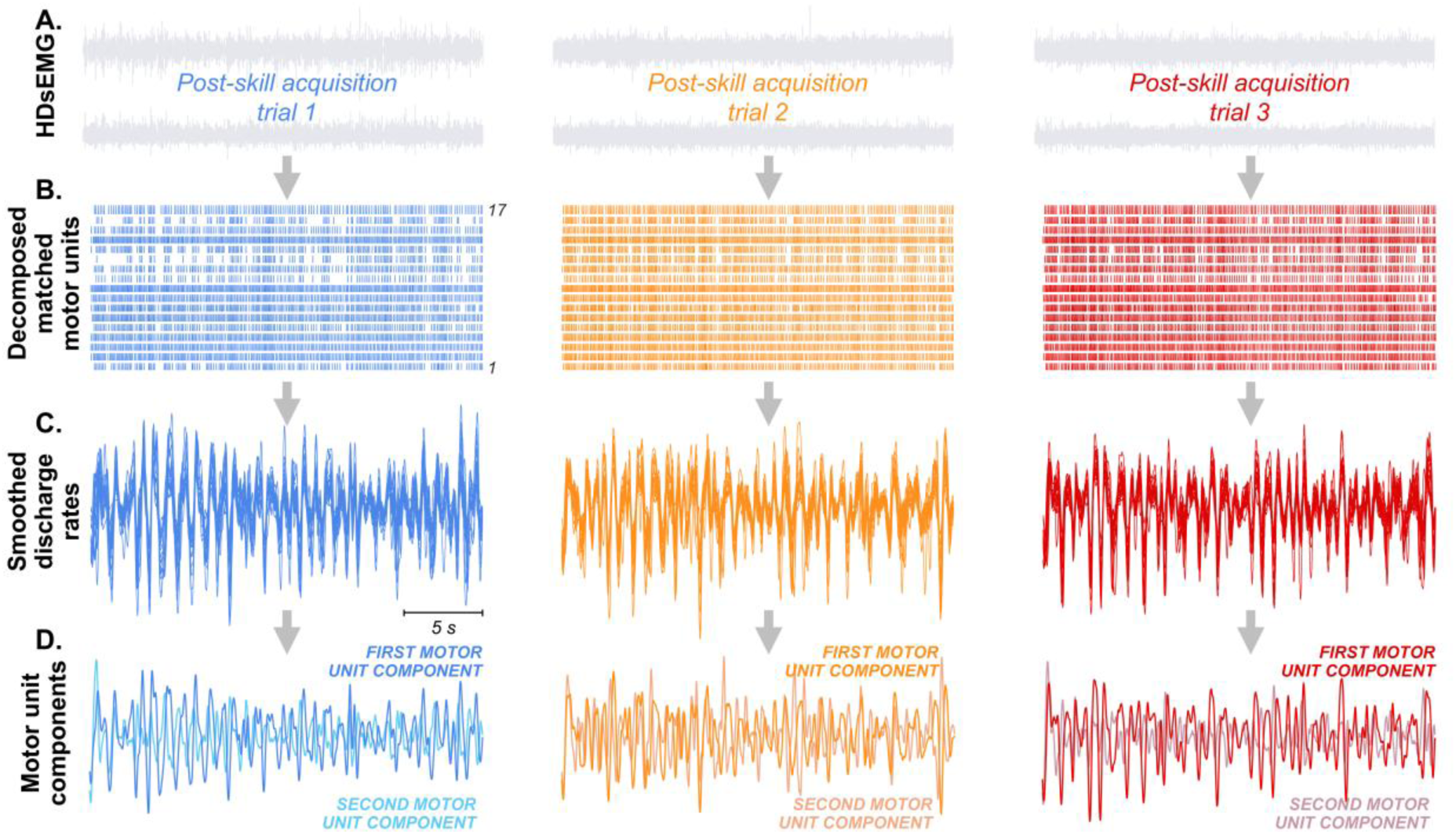
Linear dimensionality reduction analysis. For the three post-skill acquisition trials selected for analysis, high-density surface electromyograms (HDsEMG) (A) were decomposed into motor unit discharge times using a convolutive blind source separation algorithm. The motor units were tracked across trials, and their discharge times were used to calculate binary motor unit spike trains (B). Low-pass filtered discharge rates of tracked motor units were obtained by convolving the motor unit spike trains with a 400-ms Hanning window (C). The standardized and detrended smoothed discharge rates were then used to estimate low-dimensional motor unit components via principal component analysis (PCA) and factor analysis (FA). Two components were extracted for all muscles analyzed (D).

#### Calculation of smoothed discharge rate matrices

The discharge times of the motor units tracked across the three post-skill acquisition trials were used to compute binary motor unit spike trains, where a value of 1 indicates the presence of a motor unit discharge at a specific time point, and a value of 0 indicates its absence (**Figure 2B**). The low-pass filtered discharge rates of motor units were then calculated by convolving the binary motor unit spike trains with a 400-ms Hanning window (Negro et al., 2009). To remove offsets and trends, these smoothed discharge rates were high-pass filtered at 0.75 Hz using a third-order Butterworth filter (De Luca et al., 1982) and subsequently standardized to have mean 0 and standard deviation 1 (Joliffe and Morgan (1992); **Figure 2C**). The resulting standardized and detrended smoothed discharge rates, referred to as smoothed discharge rates for simplicity, were then arranged in a *r* × *c* matrix, where *r* is the number of time samples and *c* is the number of motor units. This matrix was used to estimate the neural components underlying the low-frequency oscillations of motor units by applying linear dimensionality reduction techniques, specifically PCA and FA. Additionally, a non-linear method (network-information framework) was employed to characterize motor unit networks across trials.

#### Linear dimensionality reduction techniques

Two linear dimensionality reduction techniques, PCA and FA, were used to extract the components underlying the smoothed discharge rates of motor units (Jöreskog, 1967; Joliffe and Morgan, 1992). An important step when applying PCA or FA is to determine the number of low-dimensional components to retain. Various methods have been proposed to determine this number, often based on the eigenvalues of each component (Guttman, 1954; Kaiser, 1960; Horn, 1965; Cattell, 1966; Velicer, 1976). Given previous evidence suggesting that parallel analysis is one of the most accurate methods for identifying the ideal number of components, significantly outperforming other approaches (Zwick and Velicer, 1986; Velicer et al., 2000; Hayton et al., 2004), we employed this method to determine the number of motor unit components to extract. For a detailed tutorial on performing parallel analysis, refer to Hayton et al. (2004). Briefly, we simulated a matrix of random data by shuffling the time samples of the smoothed discharge rate matrices calculated for each trial (see *Calculation of smoothed discharge rate matrices*). This random data matrix was then subjected to PCA, and the estimated eigenvalues for each component were stored. This procedure was repeated 1,000 times, resulting in a set of random eigenvalues from which the average and 95% confidence intervals were calculated. The number of motor unit components to retain was determined by identifying the eigenvalues extracted from the real smoothed discharge rate matrices that exceeded the upper bound of the 95% confidence interval of the random eigenvalues. This approach was performed separately for each of the three selected trials, and the average across trials was calculated and retained for further analysis.

Based on parallel analysis, which indicated extracting an average of one motor unit component for individual muscles and two components for the VM-VL muscles (see *Results*), we extracted two components for all analyses (**Figure 2D**). PCA and FA were applied without rotation to motor units decomposed from individual muscles (TA, FDI, and VL) as well as synergistic muscles (combined VM-VL). The VM muscle was not analyzed separately due to the low number of motor units matched across trials (see *Results*). To investigate the association between force oscillations and the two motor unit components, the cross-correlation between the detrended signals was calculated (Negro et al., 2009). To assess the consistency of the two motor unit components across trials, cross-correlation between trials was computed using 5-second non-overlapping windows, with the resulting values averaged. Additionally, the percentage of variance in the smoothed discharge rates explained by the two components obtained through PCA was calculated and retained for analysis.

#### Network-information framework

To non-linearly characterize the connectivity between motor unit smoothed discharge rates, we applied a framework combining information and network theories (i.e., the network-information framework). The detailed methodology for this approach has been described in previous studies (O’Reilly and Delis, 2022, 2024). First, the non-linear relationships between smoothed discharge rates (**Figure 3A**) were estimated using pairwise mutual information with a Gaussian copula-based approximation (Ince et al., 2017), resulting in a symmetric adjacency matrix representing the connectivities between all motor units (i.e., network-level functional connectivity; **Figure 3B**). A modified percolation analysis (Gallos et al., 2012) was then applied to determine a threshold (the percolation threshold), identifying only the significant associations between motor units (**Figure 3C**). Subsequently, graph theory was employed to construct the motor unit network, where nodes (or vertices) represent motor units, and edges (or links) denote significant associations between motor units (**Figure 3D**). We used a circular representation to visually observe the matched motor units across trials. As in the PCA and FA, this analysis was applied to motor units decomposed from individual muscles (TA, FDI, and VL) as well as synergistic muscles (combined VM-VL).

**Figure 3:**
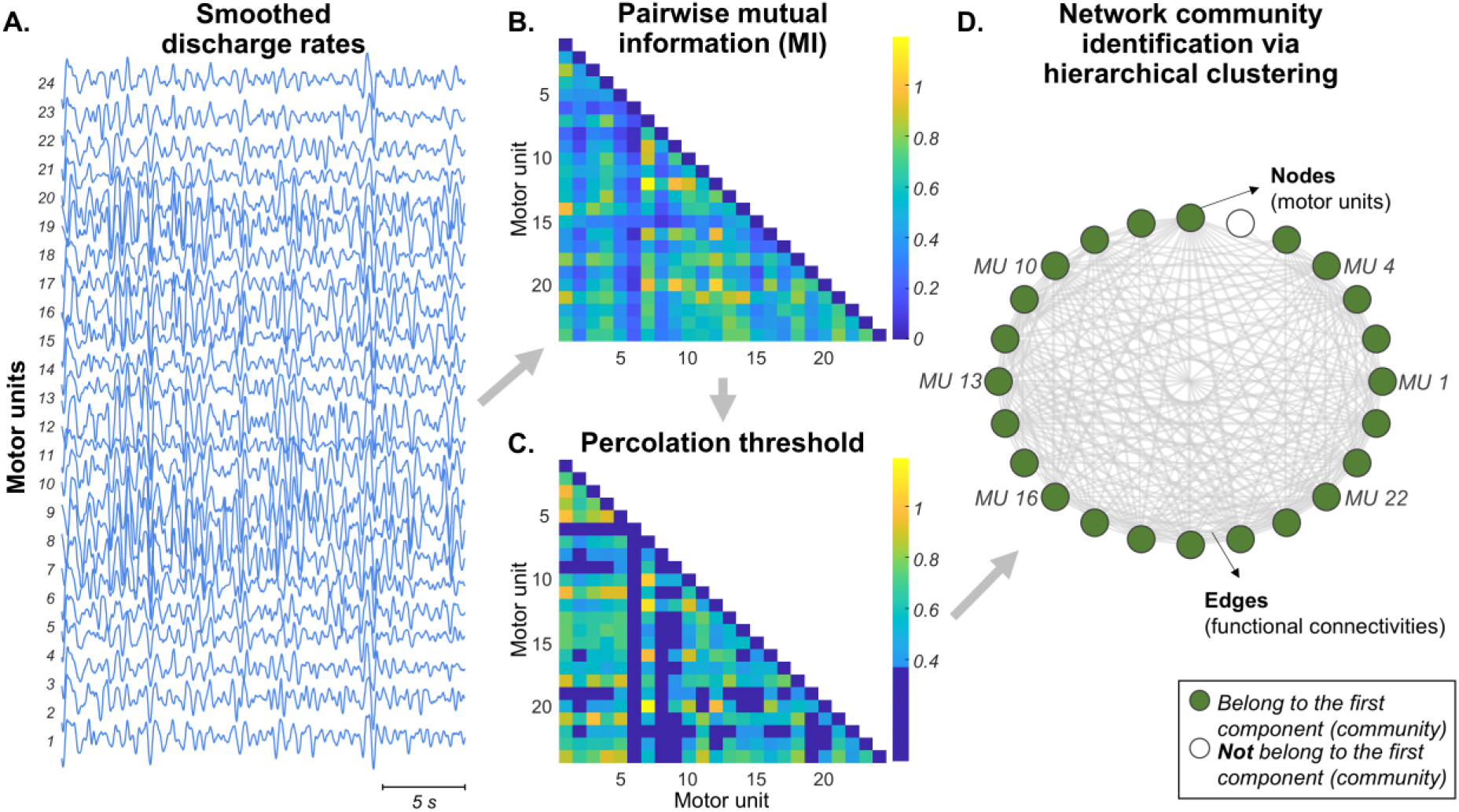
Network-information analysis. Pairwise mutual information, approximated using a Gaussian copula (Ince et al., 2017), was used to calculate non-linear relationships between smoothed discharge rates (A), resulting in a symmetric adjacency matrix (B). A modified percolation analysis (Gallos et al., 2012) identified significant associations between motor units (C). Graph theory was then applied to construct the motor unit network, where nodes (circles) represent motor units and edges (gray lines) denote significant associations (D). To identify neural components within the motor unit network, a network community detection algorithm was employed (Ahn et al., 2010), and the percentage of motor units belonging to the first component (green nodes in D) was quantified.

To identify the neural components (or communities) within the motor unit network of each post-skill acquisition trial, we applied a network community detection algorithm (Ahn et al., 2010). This algorithm quantifies overlapping components through hierarchical clustering of network links and outputs a binary matrix, where the rows represent the identified components, and the columns represent the motor units in the network. A motor unit was assigned a value of 1 if it belonged to a specific component and 0 otherwise. This process was performed separately for each trial, and the average number of identified components across the three trials was calculated. Two metrics were used to characterize the motor unit networks for each post-skill acquisition trial. First, we calculated the network density, which quantifies the number of edges in the network relative to the total possible number of edges. A higher network density indicates stronger interconnections between nodes, reflecting a more cohesive network. Additionally, we quantified the percentage of motor units belonging to the first component (represented by green nodes in **Figure 3D**).

### Statistical analysis

All statistical analyses were performed using R (version 4.3) within the RStudio environment (version 2023.06.0). To compare the RMSE between force and target, as well as the coefficient of variation of force across the three selected post-skill acquisition trials, Friedman tests were used with Bonferroni’s post hoc correction for pairwise comparisons. The Kaiser-Meyer-Olkin (KMO) measure of sampling adequacy was applied to assess whether the smoothed discharge rate matrices were factorable (Kaiser, 1974). KMO values greater than 0.70 were considered indicative of matrices appropriate for factorization (Hoelzle and J. Meyer, 2012).

Linear mixed-effect models (LMMs) were used to compare the total variance explained in the smoothed discharge rates by the first and second motor unit components. Random intercept models were applied, with trial (trial 1, trial 2, and trial 3) and motor unit component (first and second motor unit components) as fixed effects, and participant as a random effect. To compare the cross-correlation values between motor unit components and force, random intercept models were also used, with trial (trial 1, trial 2, and trial 3), motor unit component (first and second motor unit components) and linear method (PCA and FA) as fixed effects, and participant as a random effect. Similarly, to compare the cross-correlation values of motor unit components across trials, random intercept LMMs were used, with trial comparison (trial 1 vs. trial 2, trial 1 vs. trial 3, and trial 2 vs. trial 3), motor unit component (first and second motor unit components) and linear method (PCA and FA) as fixed effects, and participant as a random effect. LMMs were implemented using the package *lmerTest* (Kuznetsova et al., 2017) with the Kenward-Roger method to approximate the degrees of freedom and estimate the *p*-values. The *emmeans* package was used for multiple comparisons and to calculate estimated marginal means with 95% confidence intervals (Lenth et al., 2019).

To compare network density and the number of motor units belonging to the first component across trials, Friedman tests were performed with Bonferroni’s post hoc correction for pairwise comparisons. All individual data of motor unit discharge times recorded for each muscle in the three post-skill acquisition trials are available at https://doi.org/10.6084/m9.figshare.28324253

## Results

### Force oscillations across selected trials

From the fifteen trials performed by each participant, we selected the three consecutive trials following skill acquisition for analysis. **Figure 4A** provides a representative example of the force oscillations observed in these trials. Visually, there is a clear overlap between the force (colored traces) and target (gray traces) signals, along with a high degree of similarity in force oscillations across trials. Consistent with these observations, no significant differences in force-target RMSE were observed across trials for any of the tasks investigated (Friedman tests; *P* > 0.096 for all cases; **Figure 4B**). Similarly, no significant differences were observed in the coefficient of variation of force across trials (Friedman tests; *P* > 0.155 for all cases). For the TA, the average coefficient of variation of force values were 9.47 ± 0.42%, 9.74 ± 0.94%, and 9.73 ± 0.73% for post-skill acquisition trials 1, 2, and 3, respectively. For the FDI, they were 9.57 ± 2.28%, 9.80 ± 2.99%, and 9.30 ± 2.36% for post-skill acquisition trials 1, 2, and 3, respectively. For the VM-VL, the values were 8.65 ± 0.31%, 8.95 ± 0.27%, and 9.14 ± 0.61% for post-skill acquisition trials 1, 2, and 3, respectively.

**Figure 4:**
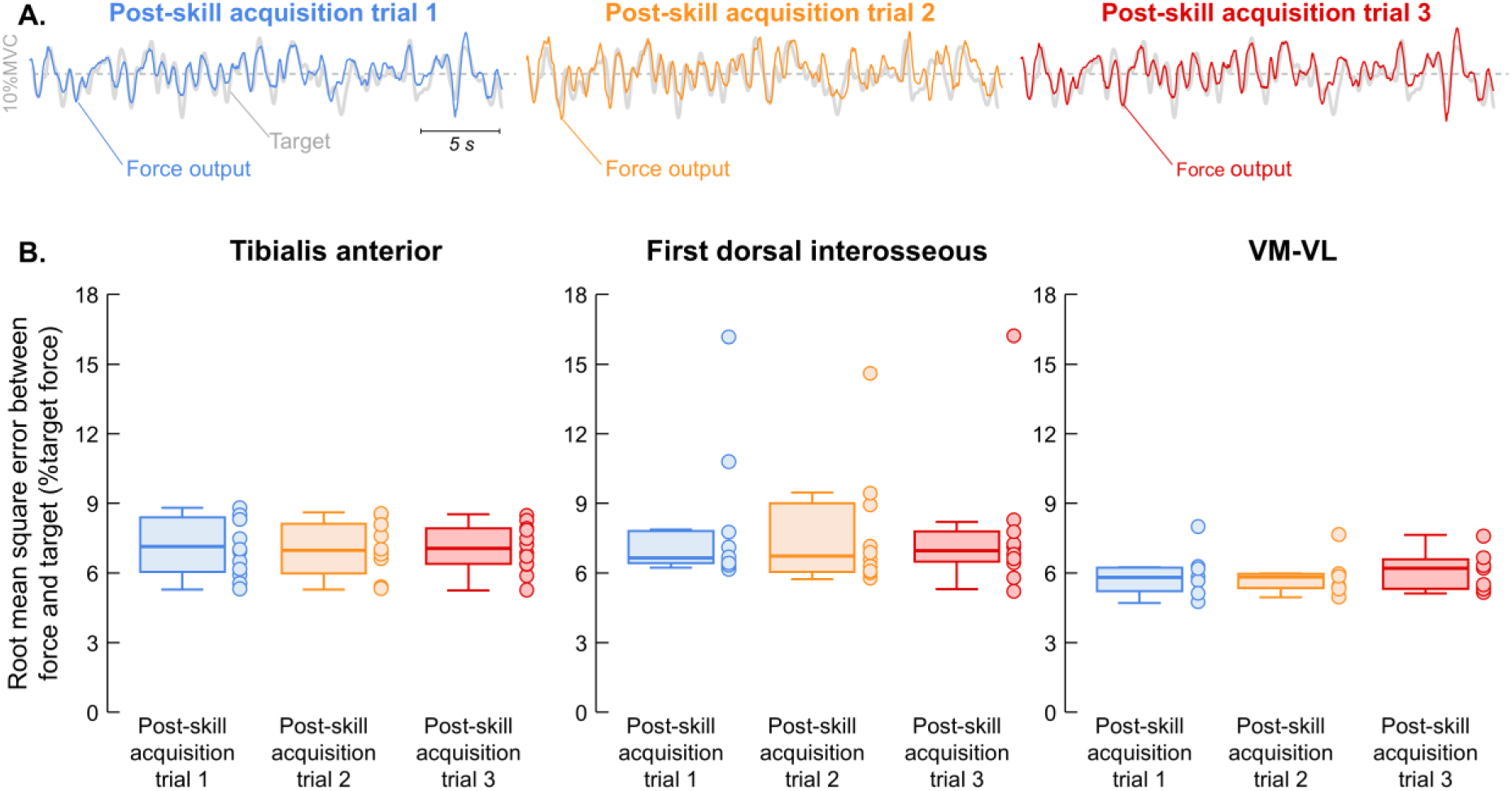
Results of force oscillations across trials. Comparison between force oscillations (colored lines) and the target (gray line) for the three consecutive post-skill acquisition trials selected for analysis (A). A high degree of similarity in force fluctuations across trials is evident. Group results of the root mean square error between the force and target are shown for the tibialis anterior (left panel in B), first dorsal interosseous (middle panel in B), and combined vastus medialis and vastus lateralis muscles (right panel in B). Circles represent individual participants. Horizontal lines, boxes, and whiskers denote the median value, interquartile range, and distribution range, respectively. VM, vastus medialis; VL, vastus lateralis.

### Characterization of low-dimensional motor unit control using PCA and FA

To evaluate low-dimensional neural control underlying motor unit activity, we decomposed HDsEMG signals into individual motor unit spike trains and tracked the motor units across trials. The average number of matched motor units per participant was 16 ± 7 for the TA, 7 ± 1 for the FDI, 3 ± 1 for the VM, and 12 ± 5 for the VL. All subsequent analysis were applied to matched motor units from individual muscles (TA, FDI, and VL) as well as synergistic muscles (combined VM-VL). We assessed whether the smoothed discharge rates of matched motor units were suitable for factor analysis. The KMO average values were 0.96 ± 0.03 for the TA, 0.90 ± 0.03 for the FDI, 0.87 ± 0.09 for the VL, and 0.91 ± 0.06 for the VM-VL, indicating that the smoothed discharge rate matrices for all muscles were appropriate for factorization.

Parallel analysis indicated that the average number of components to be extracted was 1.1 ± 0.3 for the TA motor units, 1.0 ± 0.2 for the FDI motor units, 1.6 ± 0.7 for the VL motor units, and 2.0 ± 0.7 for the combined VM-VL motor units. We opted to extract two components for all the muscles investigated using PCA and FA. For all muscles, the first motor unit component explained significantly greater variance in the smoothed discharge rates compared with the second motor unit component (LMMs; main effect of motor unit component; *P* < 0.001 for all muscles). These differences were independent of the trial analyzed (LMMs; interaction effect of motor unit component * trial; *P* > 0.412 for all muscles). The first motor unit component accounted for an average variance of 80.7 ± 6.6% for the TA, 74.5 ± 8.4% for the FDI, 55.9 ± 9.9% for the VL, and 54.3 ± 9.7% for the VM-VL. The second component explained an average variance of 4.8 ± 2.2% for the TA, 9.6 ± 3.0% for the FDI, 13.5 ± 4.2% for the VL, and 11.7 ± 3.6% for the VM-VL.

### Association between force oscillations and motor unit components

To assess how effectively the neural components underlying motor unit activity explained force oscillations, we calculated the cross-correlation between these signals. For all muscles, the first motor unit component showed significantly greater cross-correlation values with force compared to the second motor unit component (LMMs; main effect of motor unit component; *P* < 0.001 for all muscles). These differences were independent of the trial analyzed and the linear method used (LMMs; interaction effect of motor unit component * trial * linear method; *P* > 0.201 for all muscles). Specifically, for the TA, the cross-correlation values with force significantly increased from 0.07 [0.02, 0.13] for the second motor unit component to 0.52 [0.46, 0.57] for the first motor unit component (**Figure 5A**). For the FDI, the values increased from 0.22 [0.17, 0.27] to 0.56 [0.51, 0.61] between the second and first motor unit components (**Figure 5B**). For the VL, the increases from the second to the first motor unit component were from 0.06 [0.02, 0.11] to 0.50 [0.45, 0.54] (**Figure 5C**). For the VM-VL motor units, the cross-correlation increased from 0.07 [0.02, 0.13] to 0.52 [0.46, 0.57] between the second and first motor unit components (**Figure 5D**).

**Figure 5:**
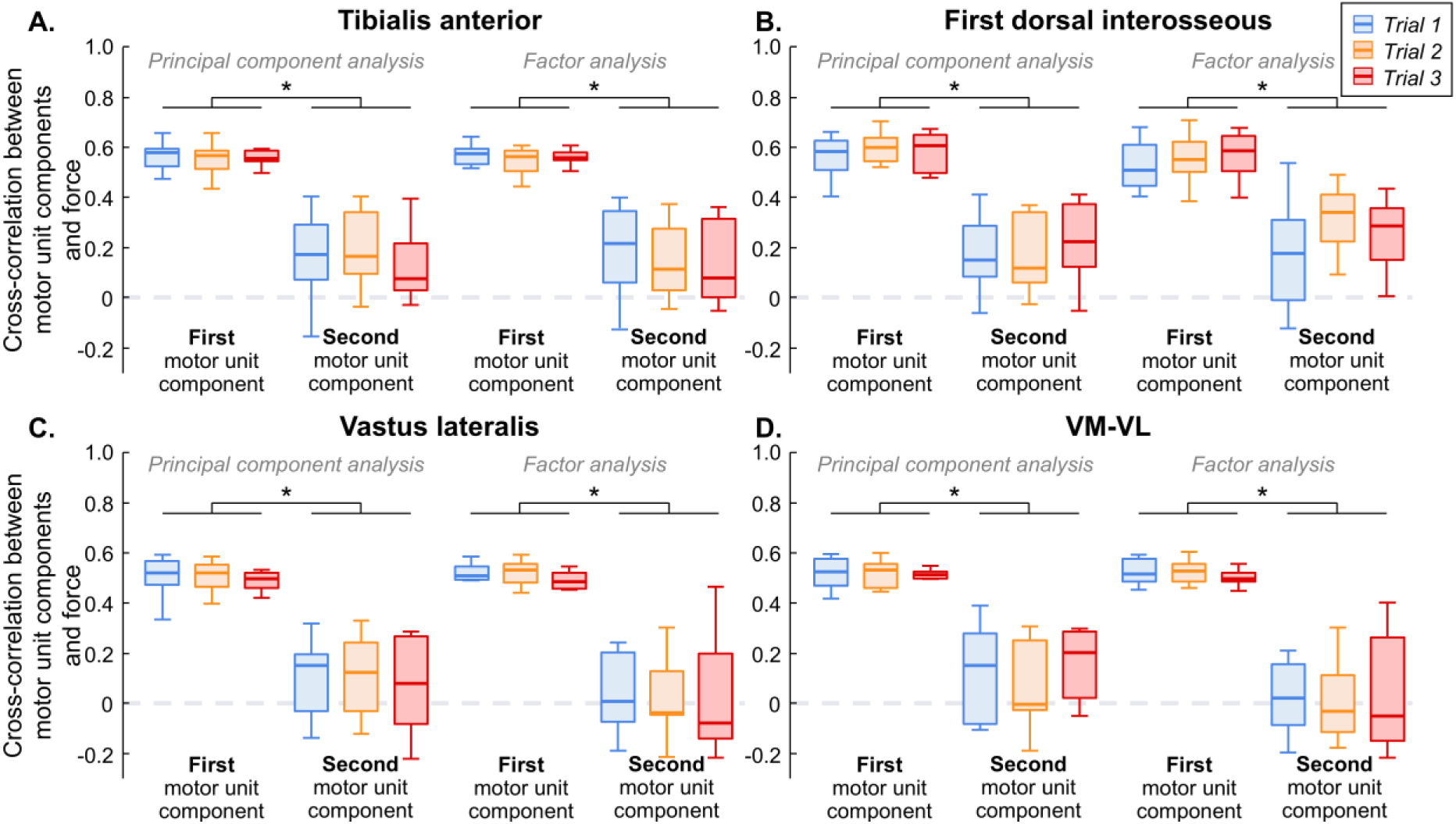
Correlation between low-dimensional motor unit components and force. To evaluate how effectively the two low-dimensional motor unit components explained force oscillations, we calculated the cross-correlation between these signals. Results are presented separately by method (principal component analysis and factor analysis) and muscle: tibialis anterior (A), first dorsal interosseous (B), vastus lateralis (C), and combined vastus lateralis and vastus medialis (D). Horizontal lines, boxes, and whiskers denote the median value, interquartile range, and distribution range, respectively. *p < 0.05. VM, vastus medialis; VL, vastus lateralis.

### Consistency of motor unit components across trials

**Figure 6** shows a representative case of the motor unit components extracted for each post-skill acquisition trial using PCA. While the first motor unit component (**Figure 6A**) remained highly consistent across trials, this similarity across trials was notably reduced for the second motor unit component (**Figure 6B**). This was in line with the cross-correlation values calculated between components, indicating an average of 0.78 ± 0.05 for the first component and 0.36 ± 0.11 for the second component.

**Figure 6:**
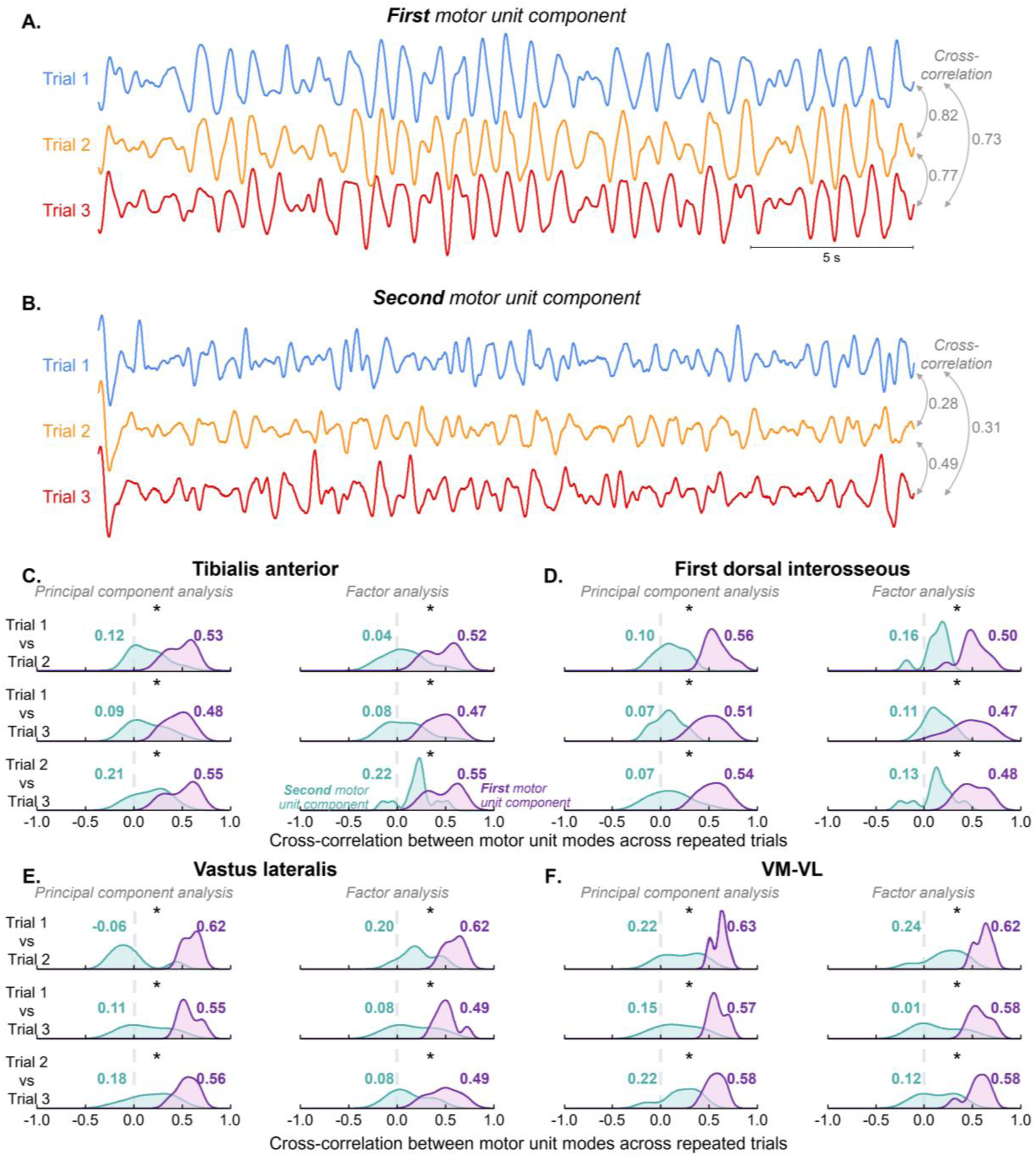
Correlation of motor unit components across trials. Representative example of the first (A) and second (B) motor unit components extracted during the three post-skill acquisition trials. Note the high similarity in oscillations of the first motor unit component across trials compared with the second component. Group results of the cross-correlation are shown for the tibialis anterior (A), first dorsal interosseous (B), vastus lateralis (C), and combined vastus lateralis and vastus medialis (D). Density curves compare the distribution of cross-correlation values for the first (purple) and second (green) motor unit components, with average values indicated. *p < 0.05. VM, vastus medialis; VL, vastus lateralis.

Consistent with the representative case, the group results revealed significantly greater cross-correlation values across trials for the first motor unit component compared to the second component (LMMs; main effect of motor unit component; *P* < 0.001 for all muscles), regardless of the trial comparison and the linear method used (LMMs; interaction effect of motor unit component * trial comparison * linear method; *P* > 0.230 for all muscles). For the TA muscle, the cross-correlation values across trials significantly increased from 0.14 [0.07, 0.21] for the second motor unit component to 0.48 [0.41, 0.55] for the first motor unit component (**Figure 6C**). For the FDI, these values increased from 0.11 [0.06, 0.16] to 0.52 [0.46, 0.57] between the second and first components (**Figure 6D**). In the VL, cross-correlation across trials increased from 0.14 [0.03, 0.24] to 0.55 [0.45, 0.66] between the second and first motor unit components (**Figure 6E**). Similarly, for the VM-VL motor units, the cross-correlation increased from 0.18 [0.08, 0.29] to 0.58 [0.48, 0.69] between the second and first components (**Figure 6F**).

### Characterization of motor unit networks across trials using network-information framework

To assess the statistical dependence between pairs of motor unit smoothed discharge rates, we characterized motor unit networks across consecutive trials using a network-information framework. **Figure 7A** shows the significant pairwise mutual information between matched motor units, along with the motor unit networks identified for the three consecutive trials. The networks showed a high degree of similarity across trials, as evidenced by consistent connections, such as those involving motor unit four (red edges) and motor unit sixteen (blue edges). Additionally, the motor unit networks were generally denser (i.e., featuring a greater number of connections), with most motor units belonging to the first motor unit component (or community), as indicated by the green circles representing these motor units.

**Figure 7:**
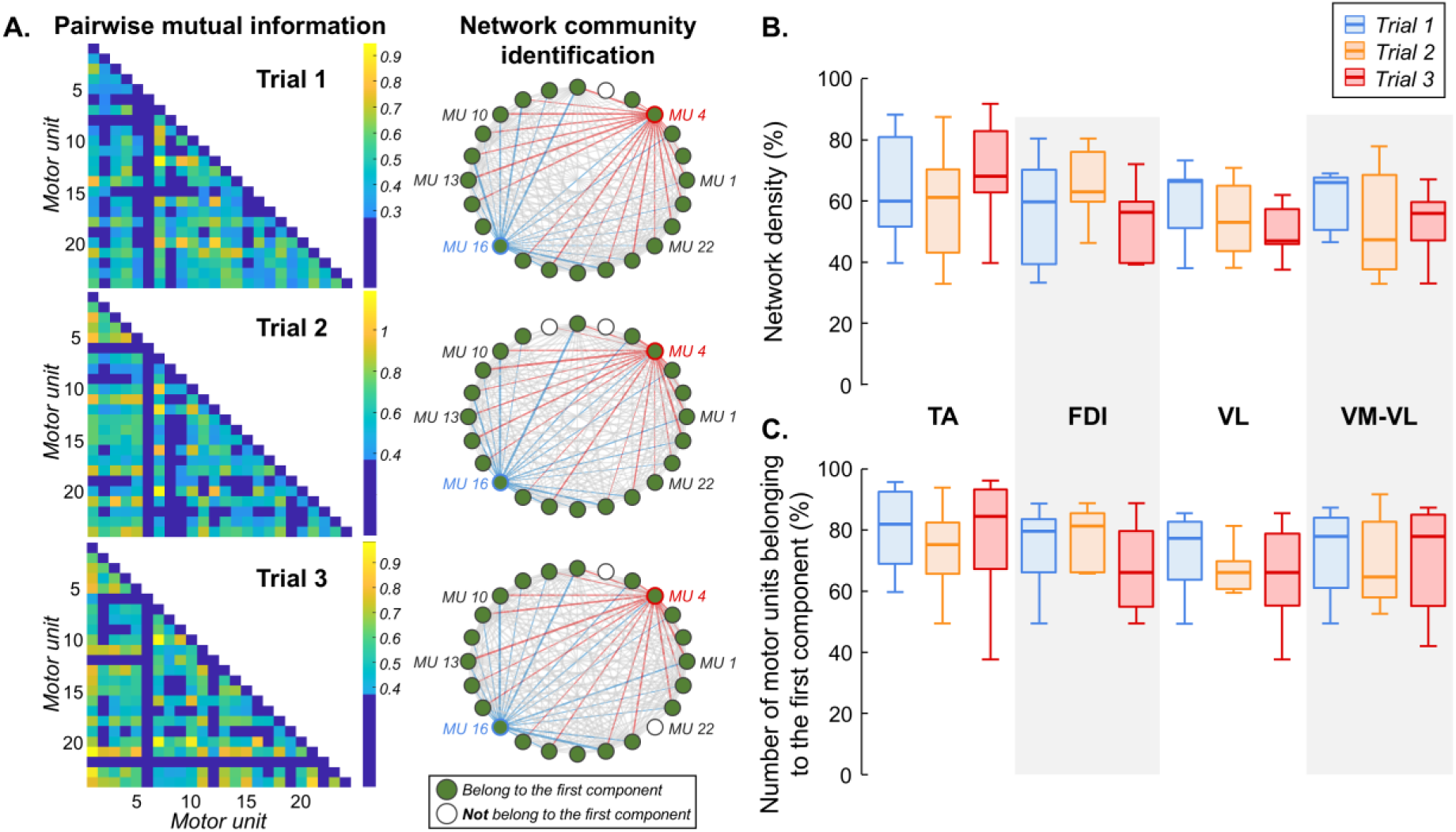
Results of motor unit networks. Representative example of non-linear pairwise mutual information analysis and the resulting motor unit networks for the three post-skill acquisition trials (A). Note the high similarity between motor unit networks, with most motor units belonging to the first component (green circles). The motor unit networks were characterized by high density (numerous connections). Group results are shown for network density (B) and the number of motor units belonging to the first component (C), separately by muscle and trial. Horizontal lines, boxes, and whiskers denote the median value, interquartile range, and distribution range, respectively. TA, tibialis anterior; FDI, first dorsal interosseous; VM, vastus medialis; VL, vastus lateralis.

For the group results, the average number of identified motor unit components in the network was 1.6 ± 1.0 for the TA motor units, 1.0 ± 0.2 for the FDI motor units, 1.3 ± 0.7 for the VL motor units, and 1.4 ± 0.6 for the combined VM-VL motor units. No significant differences in network density were observed across trials for any of the muscles investigated (Friedman test; *P* > 0.145 for all cases; **Figure 7B**). Similarly, no significant differences were found across trials in the number of motor units belonging to the first motor unit component (Friedman test; *P* > 0.345 for all cases; **Figure 7C**), except for the FDI (Friedman test; *P* = 0.047). However, pairwise comparisons with Bonferroni correction did not reveal any significant differences between trials for the FDI (*P* > 0.25 for all pairwise comparisons).

## Discussion

In this study, we investigated the relation between low-dimensional components underlying motor unit discharge rates and oscillations in muscle force during repetitive isometric tasks with highly similar force outputs. We first analyzed motor unit smoothed discharge rates of individual (TA, FDI and VL) and synergistic (VM-VL) muscles using linear methods (PCA and FA). Our results revealed that a single component was sufficient to explain most of the variance in smoothed discharge rates. Notably, the first motor unit component showed significantly greater correlations with force oscillations, compared to the second component, and remained highly consistent across trials. These findings were further supported by a non-linear method (network-information framework), which demonstrated high consistency in motor unit networks across repetitive trials, as reflected by the high network density and the inclusion of most motor units in the first motor unit component. As discussed below, these outcomes collectively suggest that during isometric contractions, motor units are primarily controlled by a single low-frequency synaptic input that closely resembles the output force oscillations.

The concept of low-dimensional neural control has been extensively studied at the muscular level, particularly through kinematics and surface EMG recordings, consistently demonstrating that many movements can be explained by a small set of weighted combinations of muscle activations (Lacquaniti et al., 2012; D’avella and Lacquaniti, 2013; Santello et al., 2013; Bruton and O’Dwyer, 2018). Earlier observations of a high degree of correlation in the discharge rates of individual motor units support the idea that low-dimensional control is not confined to the muscular level but extends to the motor unit level (Datta & Stephens, 1990; De Luca et al., 1982; Farmer et al., 1993; Kirkwood & Sears, 1978; Sears & Stagg, 1976). Common synaptic inputs across motor neuron pools are believed to be the primary source of this correlated activity and, consequently, the key determinant of alpha motor neuron control (Kirkwood and Sears, 1978; De Luca et al., 1982; Datta et al., 1991; Conway et al., 1995; Baker et al., 1997; Bräcklein et al., 2022a). While earlier evidence from this view came from correlation analysis between motor unit pairs (De Luca et al., 1982; Datta et al., 1991; Conway et al., 1995), subsequent simulation and experimental studies at the motor unit pool level demonstrated that the effective neural drive to muscles is the common synaptic input to motor neurons (Negro et al., 2009; Negro and Farina, 2011; Farina et al., 2014; Laine et al., 2015; Negro et al., 2016a; Bräcklein et al., 2022b; Rossato et al., 2024). More recent studies using dimensionality reduction methods have further corroborated the low-dimensional control of spinal motor neurons, showing that one or two neural control signals are sufficient to explain motor unit activity (Negro et al., 2009; Del Vecchio et al., 2023; Nuccio et al., 2024; Rossato et al., 2024).

Our findings align with this body of work, as we observed that a single component explained most of the variance in motor unit discharge rates during isometric tasks using linear methods (∼70% across trials and muscles). In contrast, the second component explained substantially less variance (∼9% across trials and muscles), indicating that a single component was sufficient to represent motor unit discharge patterns. These results were supported by the non-linear framework, which demonstrated highly interconnected motor unit networks, with most units belonging to the first motor unit component (**Figure 7**). The identification of a single dominant component, through both linear and non-linear methods, suggests that the CNS employs a simplified strategy for motor unit control, where common synaptic inputs play a crucial role in force control. Interestingly, while a single component was retained for individual muscles, parallel analysis indicated that an average of two components was needed for synergistic control (combined VM-VL motor units). This finding aligns with other studies on the VM and VL muscles that used different methods to determine the number of low-dimensional components to retain (Del Vecchio et al., 2023; Nuccio et al., 2024; Rossato et al., 2024). However, our results revealed that the second component accounted for only ∼11% of VM-VL motor unit activity, significantly less than the variance explained by the first component alone. Additionally, as further discussed below, only the first motor unit component in VM-VL largely explained oscillations in force (**Figure 5D**) and remained highly consistent across repeated trials with similar motor outputs (**Figure 6F**). Moreover, when applying non-linear methods, a single dominant component was identified on average in the motor unit networks, even for the VM-VL motor units. These results suggest that the number of extracted components for VM-VL motor units, particularly when using linear methods, does not necessarily reflect the true dimensionality of motor unit control. This finding is consistent with the recent study by Rossato et al. (2024), which showed that although two components were extracted from VM-VL motor units, participants were unable to dissociate motor unit pair activity during an online control paradigm. In agreement with this and prior work (Laine et al., 2015), our results support the presence of a dominant common input driving force control in the VM and VL muscles.

Mathematically, muscle force can be modeled as the convolution of the neural drive to the muscle (i.e., the cumulative motor unit spike train) with the average twitch of active motor units (Dideriksen et al., 2012; Negro and Orizio, 2017). Because twitch contractile properties act as a low-pass filter on the neural drive (Bawa and Stein, 1976; Baldissera et al., 1998; Cabral et al., 2024a), only the low-frequency oscillations in the neural drive are effectively transmitted to the force. Consequently, force oscillations are primarily determined by the low-frequency components of the common synaptic inputs to spinal motor neurons (for a review, see Farina and Negro (2015)). From a motor unit population perspective, these low-frequency components of shared inputs can be estimated from low-pass filtered motor unit discharge rates (e.g., using a Hanning window), either via dimensionality reduction techniques (Negro et al., 2009; Del Vecchio et al., 2023) or coherence analysis between motor unit spike trains (Negro and Farina, 2012; Castronovo et al., 2015; Cabral et al., 2024b; Cabral et al., 2024a). Given that dimensionality reduction methods, as applied in the current study, provide time-varying signals, their fluctuations are expected to closely resemble force oscillations if they are indeed determinants of force control. Our results corroborate this hypothesis, particularly the observation that the first motor unit component was correlated with force oscillations by ∼55% across all investigated muscles (**Figure 5**). This correlation was consistent across trials and independent of the linear method used. Notably, only the first component closely resembled force oscillations, whereas the second component showed an average correlation of just ∼12% across muscles. These findings, consistent with previous work (Negro et al., 2009), strongly suggest that a single dominant common input is the primary determinant of force control and modulation, at least for the muscles investigated in this study.

An important consideration is that, despite the strong correlation between the first component and force oscillations, the correlation values did not approach 1 (i.e., perfect correlation). Several factors could explain this discrepancy. First, motor unit discharge rates are influenced by common noise inputs, which introduce variability into the neural drive signal and affect force modulation (Harris and Wolpert, 1998; Cabral et al., 2024b). Second, the muscle-tendon system introduces non-linearities that affect the translation of motor unit activity into force output (Perreault et al., 2003; Lubel et al., 2023). Third, variability in the shape of motor unit twitches may contribute to differences between predicted and actual force oscillations. In this study, smoothed discharge rates were obtained by convolving motor unit spike trains with the same Hanning window for all motor units. Previous studies have shown that using a window shape more closely resembling motor unit twitch force can yield higher correlations between the first low-dimensional motor unit component and force fluctuations (Negro et al., 2009). Finally, while linear methods effectively capture the patterns of shared synaptic inputs (Negro and Farina, 2011, 2012), they may not fully account for the non-linear dynamics inherent in force production. Non-linear methods, such as those employed in this study (O’Reilly and Delis, 2022, 2024), can provide complementary insights by modeling the complex interactions within motor unit networks. For instance, the network-information framework revealed highly interconnected motor unit networks, emphasizing the role of shared inputs in driving force production (**Figure 7B**). Furthermore, most of the units belonged to the first motor unit component (∼80% across trials and muscles), and this pattern remained consistent across trials (**Figure 7C**). By capturing these higher-order interactions, non-linear methods can address limitations of linear approaches, offering a more comprehensive understanding of the neural control of force. The framework used in this study, along with high-order correlation methods (O’Reilly and Delis, 2024; O’Reilly et al., 2025), could be further explored in future research to provide better insights into motor unit control.

Another important finding of this study is the high consistency of the first component across trials, with average correlation values of approximately 0.6 across all investigated muscles (**Figure 6**). This suggests that low-dimensional control is not only effective in reducing the complexity of motor unit control but also provides a reliable and repeatable mechanism for force modulation. The high trial-to-trial consistency of the first component, but not the second, reinforces the idea that a single dominant control input driving motor unit activity is a purposeful strategy employed by the CNS to simplify coordination, particularly during repetitive tasks with similar motor outputs. There is an ongoing debate about whether low-dimensional control reflects a neural control scheme (i.e., hard-wired) or emerges as a consequence of task constraints (i.e., soft-assembled) (Tresch and Jarc, 2009; Kutch and Valero-Cuevas, 2012). While our experimental approach does not allow definitive conclusions regarding this question, we observed that the production and modulation of force after a new skill acquisition task was achieved through a single dominant control input that remained consistent across repeated trials of the same task.

### Methodological considerations

In this section, we would like to discuss several methodological considerations. Based on the number of trials of previous experiments involving short-term learning tasks (Knight and Kamen, 2004; Cabral et al., 2024b), our protocol was designed to ensure consistent motor outputs across trials, which was essential for analyzing the stability of motor unit low-dimensional components. Our results indicated that performing 15 trials of the proposed task was sufficient to obtain a sequence of three post-skill acquisition trials with highly similar motor outputs (**Figure 4**). This was further confirmed by the absence of significant changes in force steadiness (i.e., coefficient of variation of force) between selected trials. Compared to previous studies that examining low-dimensional control of motor unit activity (Negro et al., 2009; Del Vecchio et al., 2023; Nuccio et al., 2024), our protocol provided a distinct advantage by facilitating the evaluation of the consistency of neural control strategies.

Another important methodological point concerns the determination of the number of low-dimensional components to retain (Patil et al., 2008; Izquierdo et al., 2014; Iacobucci et al., 2022). Various methods have been proposed in the literature, including the ‘eigenvalue ≥ 1’ criterion (Guttman, 1954; Kaiser, 1960), the scree plot test (Cattell, 1966), the minimum average partials criterion (Velicer, 1976), and parallel analysis (Horn, 1965). Parallel analysis, which compares observed eigenvalues with those generated from random data, has been shown to outperform other approaches for selecting the number of components (Zwick and Velicer, 1986; Velicer et al., 2000; Hayton et al., 2004). To our knowledge, this is the first study to apply parallel analysis for selecting low-dimensional motor unit components. Given its consistency in determining the number of components across trials and muscles, as well as its data-driven nature, we recommend this approach for future studies exploring low-dimensional motor unit control.

The choice of dimensionality reduction method is another critical consideration that can affect the interpretation of results (Tresch et al., 2006; Cunningham and Yu, 2014; Bruton and O’Dwyer, 2018). While PCA and FA are often used interchangeably, they differ in theoretical and mathematical assumptions (Hoelzle and J. Meyer, 2012). PCA calculates linear combinations of measured variables (i.e., components) that maximize explained variance, including both common and unique variances. FA, on the other hand, separates common variance from unique variance, identifying latent constructs that explain shared variance among variables. Conceptually, PCA and FA differ in their approach directionality: PCA models how measured variables influence components, whereas FA assumes that latent factors drive the measured variables. For estimating commonality in motor unit discharge patterns, FA is advantageous because it focuses exclusively on shared variance across variables. However, PCA is mathematically linked to Hebbian theory (Oja, 1989, 1992; Olshausen, 1998), providing a solid foundation for its use in analyzing patterns of covariation within neuronal ensembles (Chapin and Nicolelis, 1999; Negro et al., 2009; Churchland et al., 2010; Levine et al., 2023). In this study, the choice of linear method (PCA or FA) did not affect the results for any of the investigated muscles, as we focused on correlations between motor unit components and force fluctuations, as well as consistency across similar trials. Future research should select dimensionality reduction methods a priori, based on the specific research question and underlying assumptions.

We chose not to apply component rotations in this study, as done in recent research (Del Vecchio et al., 2023; Nuccio et al., 2024), and would like to briefly address this decision. PCA assumes that extracted components are orthogonal, whereas FA can incorporate orthogonal (e.g., varimax) or oblique (e.g., promax) rotations to the initial factor solution. It is important to note that rotation redistributes variance across components, reducing the total variance explained by individual components due to changes in the partitioning of variance (Abdi, 2003). Consequently, rotating the components affects the correlations between individual motor unit discharge rates and components (i.e., loadings). For instance, oblique rotations such as promax may increase the correlation of certain original variables with one component while reducing it with another, thereby facilitating interpretation in specific applications where such associations may be expected (e.g., social sciences). In this study, we assumed uncorrelated components (i.e., orthogonal) for simplicity and conceptual clearness of the interpretation. In fact, if components are correlated, they may, in theory, be better interpreted as generated by a single synaptic input. Since component rotation is applied primarily to facilitate interpretation, future studies should investigate the physiological implications of perpendicular or oblique components.

Lastly, the non-linear analysis combining information theory and network-based approaches has provided novel insights into motor unit behavior. This framework has been recently applied to surface electromyograms to characterize neuromuscular networks (O’Reilly and Delis, 2024; O’Reilly et al., 2025). In this study, we extended this approach to motor unit discharge rates and demonstrated its potential to reveal important features of low-dimensional motor unit control during isometric contractions. By capturing complex interactions beyond traditional linear methods, this framework offers a promising tool for future research investigating motor unit network structure and its role in neural control strategies.

### Conclusions

In conclusion, our results provide strong evidence for low-dimensional neural control at the motor unit level, where a single dominant component is sufficient to explain the majority of motor unit discharge activity and force oscillations. The high consistency of this component across trials further supports the idea that low-dimensional control is a reliable and efficient strategy employed by the CNS for repetitive isometric tasks. Future studies should investigate the generalizability of these findings to other muscles and motor tasks with greater degrees of freedom. Lastly, we offer methodological recommendations to enhance the use and replicability of linear dimensionality reduction methods in motor unit research.

## Acknowledgements

This study was funded by the European Research Council Consolidator Grant INcEPTION contract no. 101045605. J Greig Inglis was supported by the Marie Skłodowska-Curie Actions Grant ‘MUDecomp’ agreement no. 101151712. Ioannis Delis was supported by the BBSRC grant no. BB/Y513799/1.

## References

Abdi H (2003) Factor rotations in factor analyses. In: Encyclopedia for Research Methods for the Social Sciences (Lewis-Beck M, Bryman A, Futing T, eds). Thousand Oaks (CA): Sage.

Ahn Y-Y, Bagrow JP, Lehmann S (2010) Link communities reveal multiscale complexity in networks. Nature 466:761–764.

Alessandro C, Delis I, Nori F, Panzeri S, Berret B (2013) Muscle synergies in neuroscience and robotics: from input-space to task-space perspectives. Frontiers in Computational Neuroscience 7.

Baker SN, Olivier E, Lemon RN (1997) Coherent oscillations in monkey motor cortex and hand muscle EMG show task-dependent modulation. J Physiol 501 (Pt 1):225–241.

Baldissera F, Cavallari P, Cerri G (1998) Motoneuronal pre-compensation for the low-pass filter characteristics of muscle. A quantitative appraisal in cat muscle units. J Physiol 511 (Pt 2):611–627.

Bawa P, Stein RB (1976) Frequency response of human soleus muscle. J Neurophysiol 39:788–793.

Bernshteĭn NA (1967) The Co-ordination and Regulation of Movements: Pergamon Press.

Bizzi E, Cheung VC, d’Avella A, Saltiel P, Tresch M (2008) Combining modules for movement. Brain Res Rev 57:125-133.

Bräcklein M, Barsakcioglu DY, Del Vecchio A, Ibáñez J, Farina D (2022a) Reading and Modulating Cortical β Bursts from Motor Unit Spiking Activity. J Neurosci 42:3611–3621.

Bräcklein M, Barsakcioglu DY, Ibáñez J, Eden J, Burdet E, Mehring C, Farina D (2022b) The control and training of single motor units in isometric tasks are constrained by a common input signal. eLife 11:e72871.

Bruton M, O’Dwyer N (2018) Synergies in coordination: a comprehensive overview of neural, computational, and behavioral approaches. J Neurophysiol 120:2761–2774.

Cabral HV, Inglis JG, Cudicio A, Cogliati M, Orizio C, Yavuz US, Negro F (2024a) Muscle contractile properties directly influence shared synaptic inputs to spinal motor neurons. The Journal of Physiology 602:2855–2872.

Cabral HV, Cudicio A, Bonardi A, Del Vecchio A, Falciati L, Orizio C, Martinez-Valdes E, Negro F (2024b) Neural Filtering of Physiological Tremor Oscillations to Spinal Motor Neurons Mediates Short-Term Acquisition of a Skill Learning Task. eneuro 11:ENEURO.0043-0024.2024.

Castronovo AM, Negro F, Conforto S, Farina D (2015) The proportion of common synaptic input to motor neurons increases with an increase in net excitatory input. J Appl Physiol (1985) 119:1337-1346.

Cattell RB (1966) The Scree Test For The Number Of Factors. Multivariate Behav Res 1:245–276.

Chapin JK, Nicolelis MA (1999) Principal component analysis of neuronal ensemble activity reveals multidimensional somatosensory representations. J Neurosci Methods 94:121–140.

Chen M, Zhou P (2016) A Novel Framework Based on FastICA for High Density Surface EMG Decomposition. IEEE Trans Neural Syst Rehabil Eng 24:117–127.

Churchland MM, Cunningham JP, Kaufman MT, Ryu SI, Shenoy KV (2010) Cortical Preparatory Activity: Representation of Movement or First Cog in a Dynamical Machine? Neuron 68:387–400.

Conway BA, Halliday DM, Farmer SF, Shahani U, Maas P, Weir AI, Rosenberg JR (1995) Synchronization between motor cortex and spinal motoneuronal pool during the performance of a maintained motor task in man. J Physiol 489 (Pt 3):917–924.

Cunningham JP, Yu BM (2014) Dimensionality reduction for large-scale neural recordings. Nature Neuroscience 17:1500–1509.

D’avella A, Lacquaniti F (2013) Control of reaching movements by muscle synergy combinations. Frontiers in Computational Neuroscience 7.

d’Avella A, Saltiel P, Bizzi E (2003) Combinations of muscle synergies in the construction of a natural motor behavior. Nature Neuroscience 6:300–308.

d’Avella A, Portone A, Lacquaniti F (2011) Superposition and modulation of muscle synergies for reaching in response to a change in target location. Journal of Neurophysiology 106:2796–2812.

Dai C, Hu X (2019) Independent component analysis based algorithms for high-density electromyogram decomposition: Systematic evaluation through simulation. Computers in Biology and Medicine 109:171–181.

Datta AK, Farmer SF, Stephens JA (1991) Central nervous pathways underlying synchronization of human motor unit firing studied during voluntary contractions. J Physiol 432:401–425.

De Luca CJ, LeFever RS, McCue MP, Xenakis AP (1982) Control scheme governing concurrently active human motor units during voluntary contractions. J Physiol 329:129–142.

De N RL (1938) ANALYSIS OF THE ACTIVITY OF THE CHAINS OF INTERNUNCIAL NEURONS. Journal of Neurophysiology 1:207–244.

Del Vecchio A, Marconi Germer C, Kinfe TM, Nuccio S, Hug F, Eskofier B, Farina D, Enoka RM (2023) The Forces Generated by Agonist Muscles during Isometric Contractions Arise from Motor Unit Synergies. J Neurosci 43:2860–2873.

Dideriksen JL, Negro F, Enoka RM, Farina D (2012) Motor unit recruitment strategies and muscle properties determine the influence of synaptic noise on force steadiness. Journal of Neurophysiology 107:3357–3369.

Ebitz RB, Hayden BY (2021) The population doctrine in cognitive neuroscience. Neuron 109:3055–3068.

Farina D, Negro F (2015) Common synaptic input to motor neurons, motor unit synchronization, and force control. Exerc Sport Sci Rev 43:23–33.

Farina D, Negro F, Dideriksen JL (2014) The effective neural drive to muscles is the common synaptic input to motor neurons. J Physiol 592:3427–3441.

Farmer SF, Bremner FD, Halliday DM, Rosenberg JR, Stephens JA (1993) The frequency content of common synaptic inputs to motoneurones studied during voluntary isometric contraction in man. J Physiol 470:127–155.

Gallego JA, Perich MG, Miller LE, Solla SA (2017) Neural Manifolds for the Control of Movement. Neuron 94:978–984.

Gallos LK, Makse HA, Sigman M (2012) A small world of weak ties provides optimal global integration of self-similar modules in functional brain networks. Proceedings of the National Academy of Sciences 109:2825–2830.

Gao P, Ganguli S (2015) On simplicity and complexity in the brave new world of large-scale neuroscience. Current Opinion in Neurobiology 32:148–155.

Gentner R, Classen J (2006) Modular Organization of Finger Movements by the Human Central Nervous System. Neuron 52:731–742.

Georgopoulos AP, Kalaska JF, Caminiti R, Massey JT (1982) On the relations between the direction of two-dimensional arm movements and cell discharge in primate motor cortex. J Neurosci 2:1527–1537.

Giszter S, Patil V, Hart C (2007) Primitives, premotor drives, and pattern generation: a combined computational and neuroethological perspective. In: Progress in Brain Research (Cisek P, Drew T, Kalaska JF, eds), pp 323-346: Elsevier.

Guttman L (1954) Some necessary conditions for common-factor analysis. Psychometrika 19:149–161.

Harris CM, Wolpert DM (1998) Signal-dependent noise determines motor planning. Nature 394:780–784.

Hatsopoulos NG, Ojakangas CL, Paninski L, Donoghue JP (1998) Information about movement direction obtained from synchronous activity of motor cortical neurons. Proceedings of the National Academy of Sciences 95:15706–15711.

Hayton JC, Allen DG, Scarpello V (2004) Factor Retention Decisions in Exploratory Factor Analysis: a Tutorial on Parallel Analysis. Organizational Research Methods 7:191–205.

Hebb DO (1949) The organization of behavior; a neuropsychological theory. Oxford, England: Wiley.

Hoelzle JB, J. Meyer G (2012) Exploratory Factor Analysis: Basics and Beyond. In: Handbook of Psychology, Second Edition.

Holobar A, Zazula D (2007) Multichannel Blind Source Separation Using Convolution Kernel Compensation. IEEE Transactions on Signal Processing 55:4487–4496.

Horn JL (1965) A RATIONALE AND TEST FOR THE NUMBER OF FACTORS IN FACTOR ANALYSIS. Psychometrika 30:179–185.

Hug F, Avrillon S, Sarcher A, Del Vecchio A, Farina D (2023) Correlation networks of spinal motor neurons that innervate lower limb muscles during a multi-joint isometric task. J Physiol 601:3201–3219.

Hug F, Avrillon S, Del Vecchio A, Casolo A, Ibanez J, Nuccio S, Rossato J, Holobar A, Farina D (2021) Analysis of motor unit spike trains estimated from high-density surface electromyography is highly reliable across operators. J Electromyogr Kinesiol 58:102548.

Iacobucci D, Ruvio A, Román S, Moon S, Herr PM (2022) How many factors in factor analysis? New insights about parallel analysis with confidence intervals. Journal of Business Research 139:1026–1043.

Ince RA, Giordano BL, Kayser C, Rousselet GA, Gross J, Schyns PG (2017) A statistical framework for neuroimaging data analysis based on mutual information estimated via a gaussian copula. Hum Brain Mapp 38:1541–1573.

Ivanenko YP, Cappellini G, Dominici N, Poppele RE, Lacquaniti F (2005) Coordination of Locomotion with Voluntary Movements in Humans. The Journal of Neuroscience 25:7238–7253.

Izquierdo I, Olea J, Abad FJ (2014) Exploratory factor analysis in validation studies: uses and recommendations. Psicothema 26:395–400.

Joliffe I, Morgan B (1992) Principal component analysis and exploratory factor analysis. Statistical Methods in Medical Research 1:69–95.

Jöreskog KG (1967) Some contributions to maximum likelihood factor analysis. Psychometrika 32:443–482.

Kaiser HF (1960) The Application of Electronic Computers to Factor Analysis. Educational and Psychological Measurement 20:141–151.

Kaiser HF (1974) An index of factorial simplicity. Psychometrika 39:31–36.

Kipke DR, Shain W, Buzsáki G, Fetz E, Henderson JM, Hetke JF, Schalk G (2008) Advanced neurotechnologies for chronic neural interfaces: new horizons and clinical opportunities. J Neurosci 28:11830–11838.

Kirkwood PA, Sears TA (1978) The synaptic connexions to intercostal motoneurones as revealed by the average common excitation potential. J Physiol 275:103–134.

Knight CA, Kamen G (2004) Enhanced motor unit rate coding with improvements in a force-matching task. Journal of Electromyography and Kinesiology 14:619–629.

Kutch JJ, Valero-Cuevas FJ (2012) Challenges and new approaches to proving the existence of muscle synergies of neural origin. PLoS Comput Biol 8:e1002434.

Kuznetsova A, Brockhoff PB, Christensen RHB (2017) lmerTest Package: Tests in Linear Mixed Effects Models. Journal of Statistical Software 82:1–26.

Lacquaniti F, Ivanenko YP, Zago M (2012) Patterned control of human locomotion. The Journal of Physiology 590:2189–2199.

Laine CM, Martinez-Valdes E, Falla D, Mayer F, Farina D (2015) Motor Neuron Pools of Synergistic Thigh Muscles Share Most of Their Synaptic Input. J Neurosci 35:12207–12216.

Latash ML, Scholz JP, Schöner G (2007) Toward a new theory of motor synergies. Motor Control 11:276–308.

Lenth R, Singmann H, Love J, Buerkner P, Herve M (2019) Package ‘emmeans’. In.

Levine J, Avrillon S, Farina D, Hug F, Pons JL (2023) Two motor neuron synergies, invariant across ankle joint angles, activate the triceps surae during plantarflexion. The Journal of Physiology 601:4337–4354.

Liddell EGT, Sherrington CS (1925) Recruitment and some other features of reflex inhibition. Proceedings of the Royal Society of London Series B, Containing Papers of a Biological Character 97:488–518.

Lubel E, Sgambato BG, Rohlen R, Ibanez J, Barsakcioglu DY, Tang MX, Farina D (2023) Non-Linearity in Motor Unit Velocity Twitch Dynamics: Implications for Ultrafast Ultrasound Source Separation. IEEE Trans Neural Syst Rehabil Eng 31:3699–3710.

Madarshahian S, Letizi J, Latash ML (2021) Synergic control of a single muscle: The example of flexor digitorum superficialis. J Physiol 599:1261–1279.

Martinez-Valdes E, Negro F, Laine CM, Falla D, Mayer F, Farina D (2017) Tracking motor units longitudinally across experimental sessions with high-density surface electromyography. J Physiol 595:1479–1496.

Muceli S, Poppendieck W, Holobar A, Gandevia S, Liebetanz D, Farina D (2022) Blind identification of the spinal cord output in humans with high-density electrode arrays implanted in muscles. Science Advances 8:eabo5040.

Muceli S, Poppendieck W, Negro F, Yoshida K, Hoffmann KP, Butler JE, Gandevia SC, Farina D (2015) Accurate and representative decoding of the neural drive to muscles in humans with multi-channel intramuscular thin-film electrodes. J Physiol 593:3789–3804.

Negro F, Farina D (2011) Linear transmission of cortical oscillations to the neural drive to muscles is mediated by common projections to populations of motoneurons in humans. J Physiol 589:629–637.

Negro F, Farina D (2012) Factors influencing the estimates of correlation between motor unit activities in humans. PLoS One 7:e44894.

Negro F, Orizio C (2017) Robust estimation of average twitch contraction forces of populations of motor units in humans. J Electromyogr Kinesiol 37:132–140.

Negro F, Holobar A, Farina D (2009) Fluctuations in isometric muscle force can be described by one linear projection of low-frequency components of motor unit discharge rates. J Physiol 587:5925–5938.

Negro F, Yavuz U, Farina D (2016a) The human motor neuron pools receive a dominant slow-varying common synaptic input. J Physiol 594:5491–5505.

Negro F, Muceli S, Castronovo AM, Holobar A, Farina D (2016b) Multi-channel intramuscular and surface EMG decomposition by convolutive blind source separation. J Neural Eng 13:026027.

Nuccio S, Germer CM, Casolo A, Borzuola R, Labanca L, Rocchi JE, Mariani PP, Felici F, Farina D, Falla D, Macaluso A, Sbriccoli P, Del Vecchio A (2024) Neuroplastic alterations in common synaptic inputs and synergistic motor unit clusters controlling the vastii muscles of individuals with ACL reconstruction. J Appl Physiol (1985) 137:835-847.

O’Reilly D, Delis I (2022) A network information theoretic framework to characterise muscle synergies in space and time. J Neural Eng 19:016031.

O’Reilly D, Delis I (2024) Dissecting muscle synergies in the task space. eLife 12:RP87651.

O’Reilly D, Shaw W, Hilt P, de Castro Aguiar R, Astill SL, Delis I (2025) Quantifying the diverse contributions of hierarchical muscle interactions to motor function. iScience 28:111613.

Oja E (1989) Neural networks, principal components, and subspaces. International journal of neural systems 1:61–68.

Oja E (1992) Principal components, minor components, and linear neural networks. Neural Networks 5:927–935.

Oliveira AS, Negro F (2021) Neural control of matched motor units during muscle shortening and lengthening at increasing velocities. J Appl Physiol (1985) 130:1798-1813.

Olshausen BA (1998) Linear Hebbian learning and PCA.

Overduin SA, d’Avella A, Roh J, Bizzi E (2008) Modulation of Muscle Synergy Recruitment in Primate Grasping. The Journal of Neuroscience 28:880-892.

Patil VH, Singh SN, Mishra S, Todd Donavan D (2008) Efficient theory development and factor retention criteria: Abandon the ‘eigenvalue greater than one’ criterion. Journal of Business Research 61:162–170.

Perreault EJ, Day SJ, Hulliger M, Heckman CJ, Sandercock TG (2003) Summation of Forces From Multiple Motor Units in the Cat Soleus Muscle. Journal of Neurophysiology 89:738–744.

Rossato J, Avrillon S, Tucker K, Farina D, Hug F (2024) The volitional control of individual motor units is constrained within low-dimensional neural manifolds by common inputs. The Journal of Neuroscience:e0702242024.

Rossato J, Tucker K, Avrillon S, Lacourpaille L, Holobar A, Hug F (2022) Less common synaptic input between muscles from the same group allows for more flexible coordination strategies during a fatiguing task. J Neurophysiol 127:421–433.

Saltiel P, Wyler-Duda K, D’Avella A, Tresch MC, Bizzi E (2001) Muscle synergies encoded within the spinal cord: evidence from focal intraspinal NMDA iontophoresis in the frog. J Neurophysiol 85:605–619.

Santello M, Baud-Bovy G, Jörntell H (2013) Neural bases of hand synergies. Frontiers in Computational Neuroscience 7.

Saxena S, Cunningham JP (2019) Towards the neural population doctrine. Current Opinion in Neurobiology 55:103–111.

Sherrington CS (1925) Remarks on some aspects of reflex inhibition. Proceedings of the Royal Society of London Series B, Containing Papers of a Biological Character 97:519–545.

Soechting JF, Lacquaniti F (1989) An assessment of the existence of muscle synergies during load perturbations and intentional movements of the human arm. Exp Brain Res 74:535–548.

Steinmetz NA et al. (2021) Neuropixels 2.0: A miniaturized high-density probe for stable, long-term brain recordings. Science 372:eabf4588.

Thompson CK, Negro F, Johnson MD, Holmes MR, McPherson LM, Powers RK, Farina D, Heckman CJ (2018) Robust and accurate decoding of motoneuron behaviour and prediction of the resulting force output. J Physiol 596:2643–2659.

Ting LH, Macpherson JM (2005) A limited set of muscle synergies for force control during a postural task. J Neurophysiol 93:609–613.

Ting LH, Chiel HJ, Trumbower RD, Allen JL, McKay JL, Hackney ME, Kesar TM (2015) Neuromechanical principles underlying movement modularity and their implications for rehabilitation. Neuron 86:38–54.

Tresch MC, Jarc A (2009) The case for and against muscle synergies. Curr Opin Neurobiol 19:601–607.

Tresch MC, Saltiel P, Bizzi E (1999) The construction of movement by the spinal cord. Nature Neuroscience 2:162–167.

Tresch MC, Cheung VCK, d’Avella A (2006) Matrix Factorization Algorithms for the Identification of Muscle Synergies: Evaluation on Simulated and Experimental Data Sets. Journal of Neurophysiology 95:2199–2212.

Tresch MC, Saltiel P, d’Avella A, Bizzi E (2002) Coordination and localization in spinal motor systems. Brain Res Brain Res Rev 40:66-79.

Velicer WF (1976) Determining the number of components from the matrix of partial correlations. Psychometrika 41:321–327.

Velicer WF, Eaton CA, Fava JL (2000) Construct Explication through Factor or Component Analysis: A Review and Evaluation of Alternative Procedures for Determining the Number of Factors or Components. In: Problems and Solutions in Human Assessment: Honoring Douglas N. Jackson at Seventy (Goffin RD, Helmes E, eds), pp 41-71. Boston, MA: Springer US.

Yuste R, Cossart R, Yaksi E (2024) Neuronal ensembles: Building blocks of neural circuits. Neuron 112:875–892.

Zwick WR, Velicer WF (1986) Comparison of five rules for determining the number of components to retain. Psychological Bulletin 99:432–442.

